# Stitching together a nm thick peptide-based semiconductor sheet using UV light

**DOI:** 10.1101/2020.07.01.183152

**Authors:** Alain Bolaño Alvarez, Marcelo Pino, Steffen B. Petersen, Gerardo Daniel Fidelio

## Abstract

Langmuir monolayer allows for two-dimensional nano-scale organisation of amphiphilic molecules. We here report that two retro-isomers peptides carefully designed to form stable monolayers showed semiconductor-like behaviour. Both exhibit the same hydrophobicity and surface stability but differ in the lateral conductivity and current-voltage due to the asymmetric peptide bond backbone orientation at the interface. The two peptides contain several tyrosines that allow for UVB-induced lateral crosslinking of tyrosines in neighboring peptide molecules. UVB-light induces changes in the lateral conductivity and current-voltage behaviour as well as monolayer heterogeneity monitored by Brewster angle microscopy that depends on the peptide bond backbone orientation and crosslinking of tyrosines. Our results indicate that one may design extended nano-sheets with particular electric properties, reminiscent of semiconductors. We propose to exploit such properties for biosensing and neural interfaces.

---

The interface between biology and electronics^1^ is ripe with numerous opportunities. Throughout molecular evolution the biological world has developed many processes that somehow is facilitating electron transport, often involving photonic input. Photosynthesis and a wide range of redox reactions are good examples. Thus, it is no surprise that soft biological material, such as peptide sheets may possess inherent electrical properties such as semiconductor properties. Biological organisms are controlled by neuronal circuits where short electrical pulses transmit information to and from organs^2,3^ helped by membrane proteins^4,5^. If we could engineer biological entities (peptides) with predefined electric properties, we may e.g. learn how to modulate or repair neuronal signalling between the brain and organs. Langmuir monolayers allow for tailor-made peptide films^6^. We describe the UVB induced synthesis of peptide-based nm-thick sheet that possess semiconductor properties. We designed two amphipathic peptides with similar surface properties but differing in its surface dipolar organization. PF1 (Ac-KKGALLLLLGYYY-NH_2_) has its hydrophilic domain at the N-terminus (Nt) whereas its retroisomer PF2 (Ac-YYYGLLLLLAGKK-NH_2_) at the C-terminus (C_t_). Peptides are orientated perpendicular to the interface with the (TYR)_3_ domain towards the aerial end and the hydrophilic (LYS)_2_ in contact with water. Both films have a high lateral stability of about 55 mN.m^−1^, similar to phospholipids, and a molecular area of 1 nm^2^.molecule^−1^. We have UVB-illuminated the peptide film in order to measure the light-induced changes in the lateral conductivity in the two retroisomers. The transversal C_t_→N_t_ and N_t_→C_t_ current-voltage was measured as well. Both peptides have conductivities 8 times higher than phospholipid film. After UVB-illumination both peptides acquired additional semiconductor properties due to the intermolecular cross linking between tyrosines in neighboring peptide molecules^7^ The conductivity and current-voltage dependence behave like a diode and they are asymmetric being greater in the C_t_→N_t_ direction compared with the opposite N_t_→C_t_, which may have functional importance for transmembrane proteins. We postulate that the asymmetric peptide bond backbone determines the peculiar conductivity and the current-voltage dependence behaviour. We therefore conclude that we can design peptide-sheets with predefined chemical as well as semiconductor properties. Peptides may therefore adopt active electronic roles in biosensors with the possibility that the novel properties may increase the sensitivity of the biosensors.

Since the Nobel Prize in Chemistry in 2000 for the discovery of conductive organic polymers^8^ has emerged the interest of about the electronic semiconductor properties in peptides^9^. Since late 1950s many investigators have been contributed to describe the electron transfer machinery in cells^10^. Beratan et al. described a model of electronic coupling in proteins, which was named as *Tunneling Pathway*^11^. These authors framed the interactions in a folded polypeptide to three types: covalent bonds, hydrogen bonds, and through-space contacts^11^ such as van der Waals contact between atoms^12^.

2D peptide nanofilms could potentially have applications as an organic conducting films, in biocatalysts and in bioelectronics^13-16^. Others authors have also reported that microbial nanowires and peptides as potential candidates for bioelectronics material^17^. Also, some authors have claimed that these studies would help us to further understand the transduction signal phenomenon in proteins^18^.

In the present work, we present a *2D* nano sheet conductivity and current-voltage studies by using two identical amphiphilic peptides with reversing sequences that differs in its surface dipolar organization with similar surface properties. The amphiphilicity of both peptides is asymmetric. Both peptides are acetylated and amidated at the Nt or Ct, respectively (Fig. 1). Additionally, we show that the semiconductor properties acquired by both peptides are modified after UV illumination due to the intermolecular di-Tyr and iso di-Tyr formation.

**Fig. 1.**
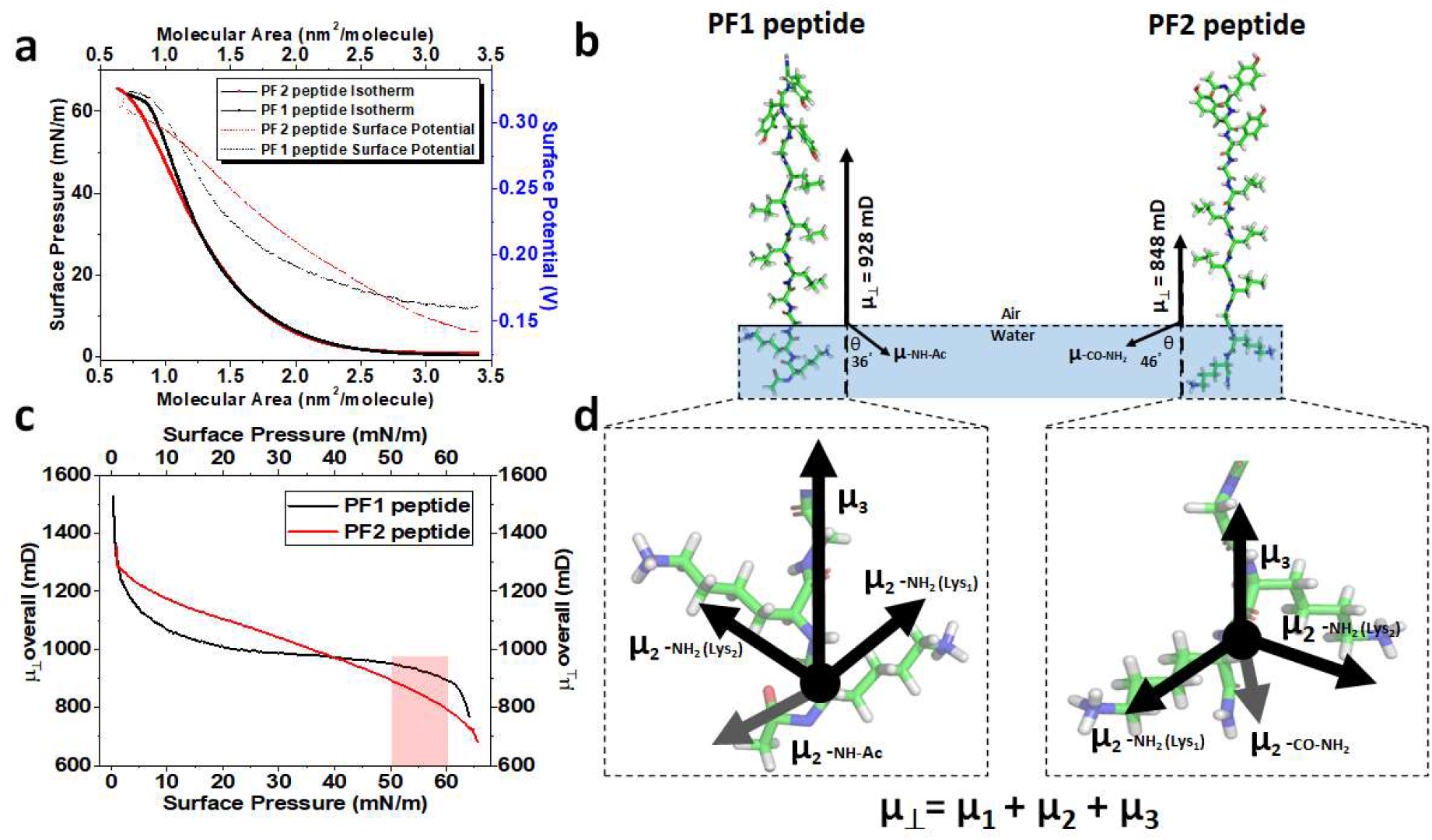
Surface properties and dipolar organization of PF tyrosinated peptides at the air-water interface. **a**, Surface pressure-area (π-Area, in straight line) and surface potential (in dot line) isotherms for PF1 (in red) and PF2 (in black) peptides. **b**, Stick representation of both peptides with the most probable conformation: extended β-sheet perpendicular at the air-water interface (using Pymol software) with the overall dipole moment (μ_⊥_) and the displacement angle of the hydrophilic portion from low packing (5 mN.m^−1^) to high packing (55 mN.m^−1^). **c**, Overall dipolar moment (μ_⊥_) dependence of PF1 and PF2 with surface pressure. **d,** Zoom in of the structural representation of the hydrophilic portion for both peptides with the 3D projection of the μ_2_ dipole contributions.

## Surface properties and dipolar organization of tyrosinated amphiphilic peptides

Our results have shown that both PF peptides behave a lipid-like behaviour at air water interface (Fig.1). The π-A isotherms show a typical profile similar to those observed for hydrophobic signal sequence peptide^19^. Both peptides have a high stability with a collapsing point of around of 55 mN.m^−1^ and limiting molecular area (at high packing) of around 1.12 nm^2^. molecule^−1^ (Fig. 1A). The perpendicular cross-section of this molecular area is compatible with a β-extended structure perpendicular at the air-water interface^19^. In this interfacial organization, each peptide is in contact with the water through the LYS residues corresponding to the hydrophilic end portion (equivalent to the polar head group of a lipid).

The sequence reversion in an extended β-sheet peptide does not impose major differences in the overall dipole moment probable due to the a more parallel orientation of the peptide bond with respect to interface^19^. Thus, as expected, the surface potential and overall dipolar moment found for both peptides only display minor differences, with some particular behaviour upon compression, (see Fig. 1). The analysis of the dipolar moment of PF2 compared to PF1 shows an experimentally difference in dipolar organization at the interface at high packing, close to 55 mN.m^−1^ (Fig. 1C). Taking into account that the values of surface potential ΔV of both films are positive, the overall dipolar moment (μ_⊥_) towards the air end for an ionized monolayer (according to^20,21^) is given by: 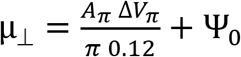 where A_π_ and ΔV_π_ are the molecular area and the surface potential at a defined surface pressure π, in nm^2^ and millivolt units, respectively; Ψ_0_ is the potential difference of the ionic double layer of the ionized peptide film (Ψ_0_ values are practically the same because of the surface charge density is identical for both peptides). μ_⊥_ is the overall dipole moment perpendicular to the air/water interface resultant of three contribution according to μ_⊥_= μ_1_ + μ_2_ + μ_3_ (see^21,22^). μ_1_ comes from the oriented water molecules around the hydrophilic portion of the amphiphilic peptide; μ_2_ is the contribution of hydrophilic end corresponding to the lysine residues (equivalent to the polar head group in the lipids) and μ_3_ comes from the hydrophobic portion of the peptide corresponding to the leucine and tyrosine domains (equivalent to hydrocarbon chains in lipids), see Fig. 1d.

At low packing, around 5-10 mN.m^−1^, the overall μ_⊥_ of PF2 is more positive than PF1 and the value is reversed at high packing, see Fig. 1c. This difference is probably due to a higher dependence of μ_1_+μ_2_ with packing for PF2 in relation to PF1. Accordingly to the theoretical interpretation of surface potential in monolayers^20^, the average dipolar moment μ_⊥_ is due to the effective perpendicular moment 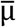, making some angle θ with respect to the vertical to the interface, i.e. that 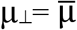 cos θ. Assuming a close perpendicular orientation of both peptides at the interface is possible to calculate a relative angle of tilting assuming the orientation at high packing with a cos θ = 1. The change in the angle orientation of μ_1_+μ_2_ contribution (the lysines and the amide group given by the domain −GKK-NH_2_ plus the hydration water) for PF2 is around 46° when it is compressed from 5 mN.m^−1^ to 55 mN.m^−1^, whereas the value for PF1 with its hydrophilic domain Ac-KKG-is about 36°. This is indicating a more rigid hydrophilic portion for PF1 compared with PF2 (or less compression dependence). We commented other chemical details in Supplemental material (see, Fig. S1).

## Surface cross-link formation of tyrosinated peptides upon UV illumination

After peptide spreading, both tyrosine-rich amphiphilic compounds were exposed to UVB illumination, by a period of time, by using a UV LED lamp (see Fig. S2 for details). At regular intervals the formation of di-Tyr was followed by monitoring the fluorescence emission at 410-420 nm^23,24^. We have also used RAMAN microscopy to follow di-Tyr and iso-di-Tyr formation (see Fig. S3 for details).

## Post-illuminated tyrosinated peptides based nano thick sheet

We coupled our UV illumination source setting to a Brewster Angle Microscopy (BAM) to investigate if the illumination and di-Tyr formation at the surface induces interfacial heterogeneity. We performed BAM before and after UV illumination taken the images at interval of 10 minutes. We observed heterogeneity after illumination in both peptides. However, PF1 shows detectable more roughness than PF2 (see Fig. 2). The heterogeneity is deducted from the reflectivity values of the films upon illumination. The observed heterogeneity for both peptides after illumination is attributable to changes in the surface organization of the film due to the cross-linking of tyr domains in PF2 peptide and, also di-iso-di-Tyr formation in PF1 peptide (see Fig. S3 b). We estimate the heterogeneity by using the standard deviation ratio media comparing the brightness intensity throughout all the image observed in the BAM (see Material and Methods for details). In comparison, PF1 shows more heterogeneity than PF2 after one-hour of UV illumination (Fig. 2a and 2b). The greater heterogeneity in PF1 peptide may be related to the coexistence of different types of cross-linking (di-tyrosine and iso-di-tyrosine) compatible with the different pattern obtained for RAMAN and fluorescence spectroscopy (compare Fig. S3). Thus, we show that it can be designed peptide-based sheets with predefined chemical and peculiar semiconductor properties (discussed below).

**Figure 2:**
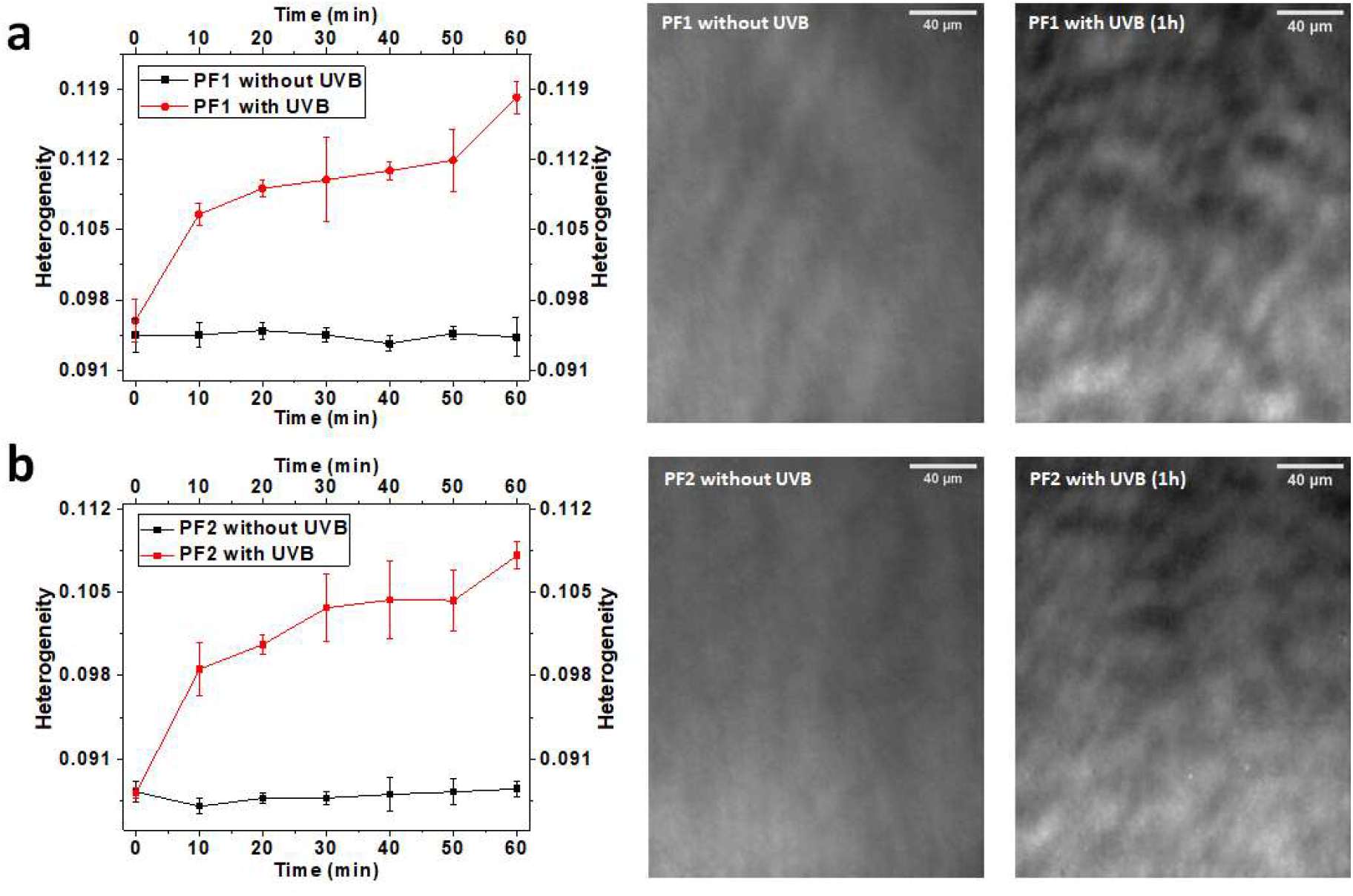
Heterogeneity of tyrosinated peptide films upon illumination to form a semiconductor sheet. **a,** Heterogeneity analysis of the taken images using BAM for non-illuminated film (black) and UV-illuminated film (red) for PF1. **b,** Heterogeneity analysis of the taken images using BAM for non-illuminated film (black) and UV-illuminated film (red) for PF2. BAM image for PF1 (right upper part) or PF2 (left lower part) after one hour of UV illumination. Have a note that PF1 is more heterogeneous than PF2. The BAM images contrast were enhanced for a better visualization.

## Semiconductor properties of tyrosinated peptides

We have designed three electronic devices for: ***d1)*** to switch on-off the UVB lamp to illuminate the peptide film to induce di-Tyr formation, ***d2)*** to measure film ***lateral conductivity*** and ***d3)*** to measure ***the transversal C_t_→N_t_ and N_t_→C_t_ current-voltage dependence*** of the spread peptide film on the air/water interface. Also, we coupled to this device the UVB illumination source allowing us the simultaneous measuring of conductivity upon illumination. The UVB light over the negatively polarized plate (polarization) down the tyrosine rich peptide monolayer induces a photoelectric effect that can be measured by the changes in the resistance of peptide releasing electrons thorough out the monolayer.

In all conditions we observed a higher conductivity for PF2 compared with PF1 peptide. Some authors had proposed that an increase in the electrical conductivity of the delocalized electrons in aromatic amino acids when they dynamically interact through near side chains causing hopping charge transfer^25,26^ which could explain that the lateral electron flow (conductivity) is higher in C_t_ to N_t_ (water to air) orientation of PF2 peptide than N_t_ to C_t_ (water to air) orientation of PF1 peptide when polarization and ***d1 off*** mode, (see Fig. 3b blue box and 3c blue curve).

**Figure 3:**
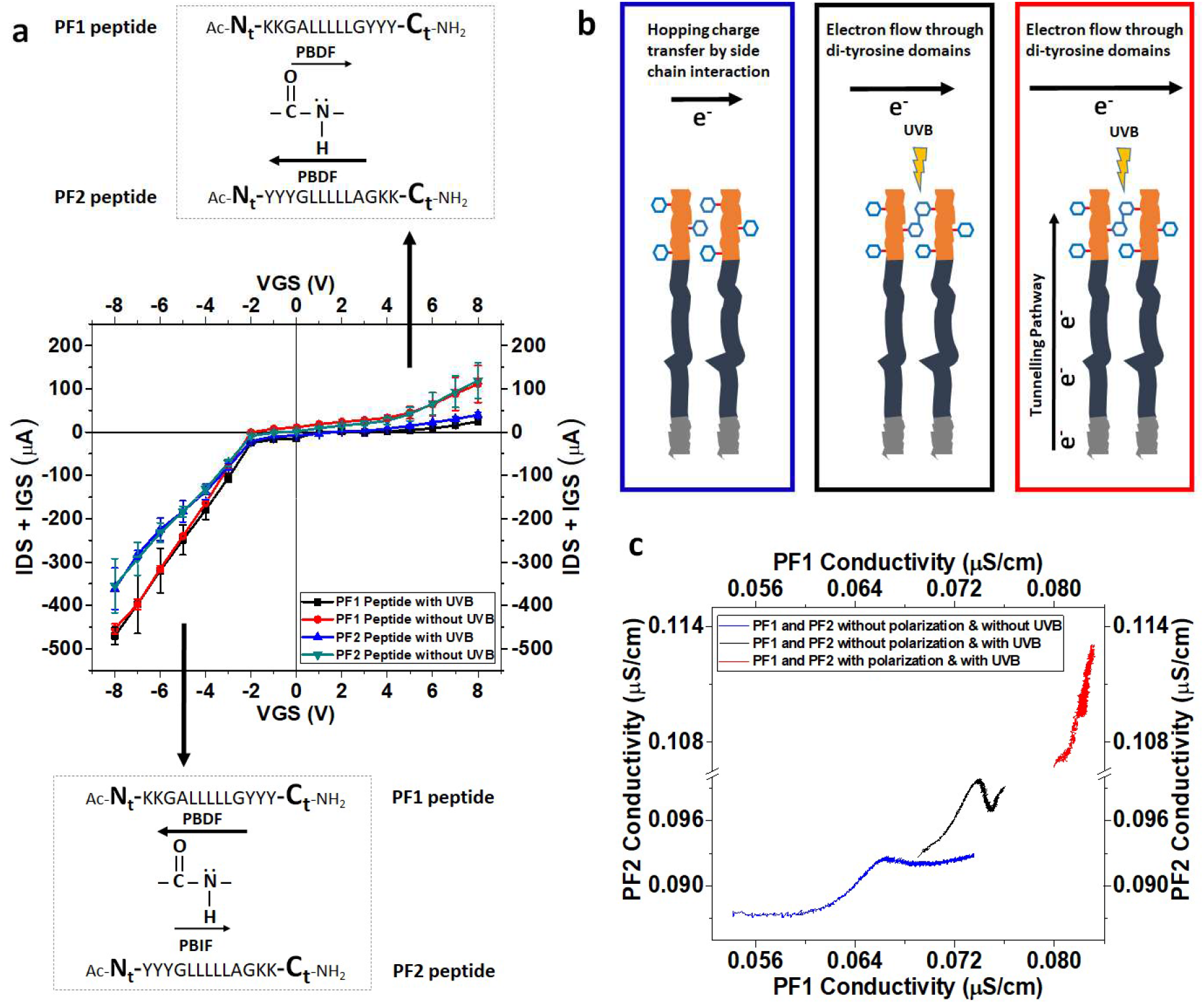
Conductivity and current-voltage dependence of PF1 and PF2 peptides. **a**, Peptide diode behaviour representation with their respective directions and relative magnitudes of the current electron flow through the oriented tyrosinated peptide backbone from N_t_ to C_t_ for PF1 (water to air) and from C_t_ to N_t_ for PF2 (water to air). **b**, Schematic diagram of experimental conditions during conductivity measurements, **(blue)** polarization and UVB light in switch off mode, the electron flow is by hopping charge transfer by side chain interaction^25^; **(black)** polarization switch off and UVB light in switch on mode, the electron flow throughout the di-Tyr domains; **(red)** polarization and UV light in switch on mode, electron flow throughout the peptide backbone by tunnelling covalent pathway^11^ summed to electron flow throughout the di-Tyr domains. **c**, relationship of conductivity of PF2 vs. PF1 peptide ratio according to the conditions displayed in **b** (note that PF2 peptide conductivity is higher than PF1 peptide in all experimental conditions). This relationship is built from the ratio of conductivity vs time showed in Fig. S4. The acronyms **PBDF** and **PBIF** mean the Peptide Bound Direct Flow and Peptide Bound Inverse Flow, respectively; see the text. For a schematic draw of the circuit to measure conductivity and current-voltage dependence refer to Fig. S4.

When ***d1*** is ***on*** and the polarization is in ***off*** mode (see black box in Fig. 3b), the relative lateral conductivity of PF2 is increased with respect to PF1 (compare black box with the blue curve in Fig. 3c). This effect is due to the augmented interchain conductivity through the di-Tyr formation upon UV-illumination. In the third condition, when ***d1*** and polarization are in ***on*** mode (see Fig. 3b red box and 3c red curve) besides the lateral interchain conductivity throughout the di-Tyr domain, we observed an additional intrachain conductivity due to the peptide backbone by tunnelling pathway^11^, the scores are higher for PF2 than PF1 (red box representation in Fig. 3b and red curve in Fig. 3c).

A fact to be highlighted that pure lipid monolayer (DMPC) has less conductivity when compared with both PF1 and PF2 peptides. So, lipids not only impose a higher resistance barrier across the membrane (in transversal way) but also impose a very low conductivity ***in the lateral electron current movement*** (see Fig S4). The non-tyrosinated KL-LK retro isomer peptides have similar lateral conductivities properties regardless of the isomerism (molecular asymmetry) (see Fig S4). Thus, it would appear that the inclusion of a tyrosine domain far away from the hydrophilic portion in the C_t_ → N_t_ backbone orientation facilitates the electron flow.

Thus, the sequence reversion does not affect the surface properties (lateral stability) but it is dramatically affecting the conductivity (and semiconductor) properties and the diode-like behaviour as it is mentioned below.

We measure the current by scanning a range of voltage polarization from −8 V to +8 V. Thus, we measure the changes in I_D-S_ (drain to source current) + I_G-S_ (gate to source current) under desired voltage polarization after 5 hour of UV illumination hopping for the highest degree of tyrosine cross linking, see Fig S4. The results show that both peptides PF1 and PF2 have a diode-like behaviour.

We observe a different behaviour in the current flow in illuminated interface (with di-Tyr formation) respect to the non-illuminated condition. Illuminated peptides, show higher resistance to the current flow than non-illuminated condition (Fig. 3a upper panel). However, as we show in Fig. 3a lower panel, the resistance to current flow was similar in both conditions (illuminated vs. non-illuminated), but it is noteworthy that PF1 showed more absolute current than PF2. It is clear that the formation of di-Tyr increases the diode-like behaviour in both peptides (see Fig. 3a, upper panel). For comparison, we have used clean air-water interface and non-tyrosinated KL and LK retro peptides as controls^6^. In all cases, the behaviour of non-tyrosinated peptides shown a more resistance property compares with the tyrosinated ones (Fig. S4).

Others authors have said that the paths of the electron transfer process are localized in the backbone^27^ due to the peptide bond, nitrogen (**N**) is the donor^28^, whereas oxygen (**O**) from the carbonyl group is the acceptor with a higher electronegativity^29,30^ determining polarity in the peptide bond and creating a way for electron flow through the backbone. So, it is expected that the electrons flow is favoured from **N** to **O** which is assimilable to the **C_t_** to **N_t_** direction. We named this direction of electrons flow as “Peptide Bound Direct Flow” (**PBDF**). Thus, the less favourable electrons flow direction from **O** to **N** is assimilable to **Nt** to **C**t direction. This less favourable electrons flow direction from **O** to **N** is defined as “Peptide Bound Inverse Flow” (**PBIF**). In PF1, from tyrosines to lysines domain, the electrons flow is favourable but, the reverse, from the lysines to tyrosines is unfavourable. Thus, the inverse situation found for PF2 (Fig. 3a).

In this connection, some authors shown that the electron transfer rates depend on the direction along the peptide backbone^31^, specifically it is faster from **C_t_** to **N_t_** terminus than the opposite direction^32^. However, we presented two reverse peptides with a β-Sheet conformation with its individual dipole moment in the same direction and with close values (Fig 1b) indicating that for electron flow is more important the orientation of the peptide backbone that the overall dipole moment. However, in the surface the structural orientation of the peptide backbone in α-helix is different^19^ and it was reported that the dipole moment has a major influence on the electron transfer^33–35^ accelerating it in the same direction of the α-helix in the peptide backbone^33^. Thus, the conduction from **C_t_** to **N_t_** is faster than **N_t_** to **C_t_** (Fig. 3a) similarly to the data reported using α-helical polyalanine monolayers^32^ in accordance with Page et al. have shown that the electron transfer process in proteins is given regardless of the conformation and heterogeneity^27^. However, it is legitimate to consider that there are subtle differences ***in the behavior of electron flow***. Indeed, we have observed changes in the conductivity of oriented tyrosinated peptides, either the direction is **N_t_→C_t_** or **C_t_→N_t_**. In this connection, but with a different approach, it was recently reported that the conductance in protein-ligand systems is sensitive to the charge and the nature of chemical electrical contact between them^36,37^. On the other hand, we constructed a conceptual assimilation of peptide bond backbone to an inorganic semiconductor with diode behavior (see Fig S4 and additional text in Supplementary Material).

## Significance of the findings

Taking into account that other authors point out there is not currently known biological role of electronic conductance in proteins^37^, extrapolating our experimental data to proteins embedded in lipid bilayers regarding to information flow across of membranes we hypothesized its intrinsic asymmetry. Fig. 4 shows a cartoon representation of a hypothetical polytopic transmembrane protein with its N-terminus inside and its C-terminus out. As we have shown, electronic information is different if it is considered that the electrons have less resistance to flow when it goes from **C_t_** to **N_t_** compared to the reverse flow from **N_t_** to **C_t_** given by its direction peptide backbone (Fig. 3a). This peculiar behavior conducted us to give an additional conceptual framework of spanning transmembrane proteins. So, each transmembrane peptide not only is given structural stability as being part of the membrane in a high amphipathic environment with restriction imposed by the lipid bilayer but different to the lipids, it also gives asymmetric in the in-out electronic information that can be fluxed throughout them.

**Figure 4:**
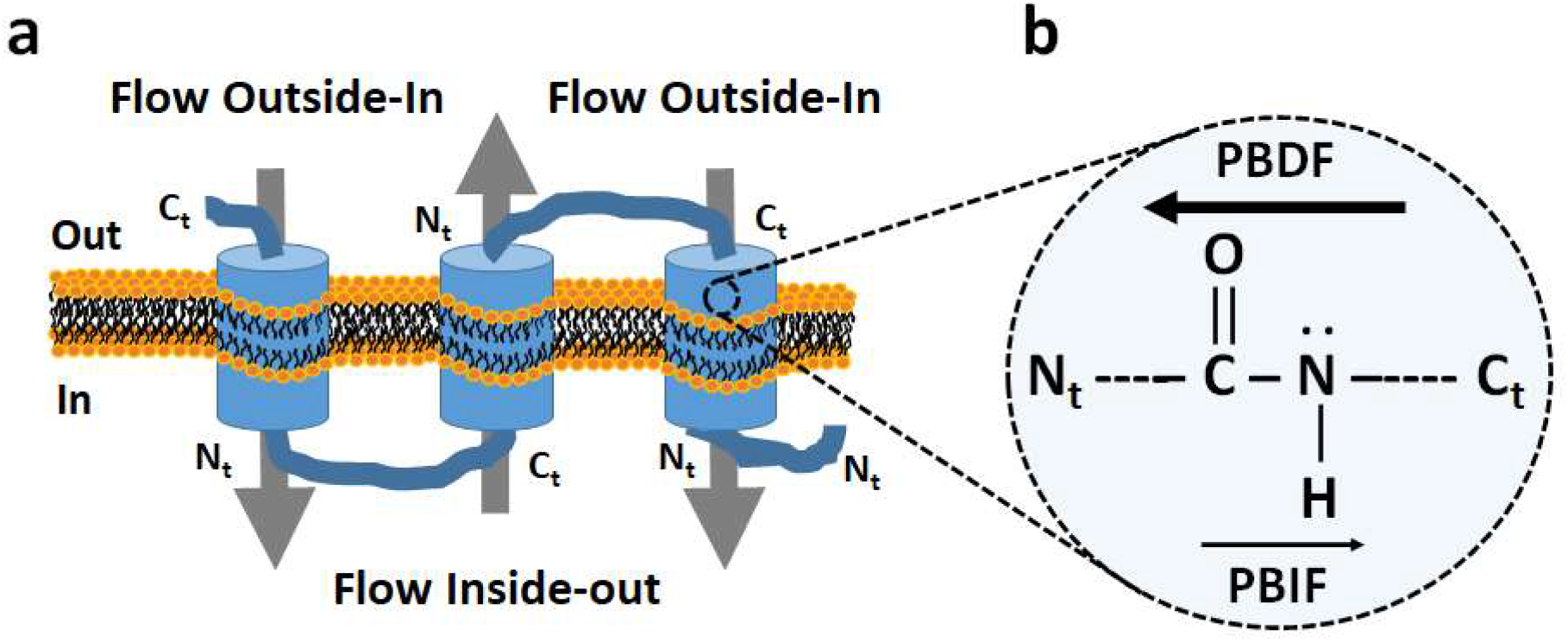
Cartoon representation of a hypothesized asymmetric transmembrane flow information. **a**, Asymmetric representation of a (multi)spanning transmembrane protein domain showing the different in-out bi-directional transmembrane information flow. b, Zoom in of the transmembrane segment showing the peptide bond direction flow. PBDF, peptide bond direct flow; PBIF, peptide bond inverse flow.

The bidirectional signaling across membrane, from a structural point of view, it has been postulated to occur for integrin proteins.^38^ In this connection, it is postulated that there are mechanisms in the membrane that can operate in both: the inside-out and outside-in directions. On the case of integrins, signaling across the plasma membrane is coupled to extracellular conformational changes leading to the separation of transmembrane domains^38^. Additionally, for membrane proteins with spanning transmembrane segments the asymmetric electronic information occurs because each **C_t_→N_t_** segment is alternatively located in the membrane following a **N_t_→C_t_** segment, see (Fig. 4). The increased conductivity due to 265 nm UV light exposure may have applications in optogenetics^39^. The therapeutic treatments for neurological disorders is limited due of the lack of scalable biomaterials at molecular level for neuronal systems that allow functionalization of cell surface with specific bimolecular array for repairing damage in some specific neurons with minimal immune response^40^.

The design of quantum dot-peptide bioconjugate systems for monitoring proteolytic activity using fluorescence resonance energy transfer (FRET) has been reported elsewhere. The authors did not mention that the conjugated peptide is oriented from Nt to Ct linking donor to acceptor (the less efficient orientation for conductance). However, they do emphasis that the FRET efficiency improves as the number of peptides assembled to quantum dots increased^41^. Keeping in mind our results, the FRET efficiency could be improved with the inverse peptide sequence linker (from C_t_ to N_t_, the orientation with higher efficiency for conductance) and it would be necessary less linking peptide acceptors coupled to the quantum dot. On the other hand, the conductivity behavior of organized amphiphilic tyrosinated peptides upon illumination may be suggesting a peptides sheet (Fig. 2) based substrate for the construction of bioinspired organic solar cells instead of organic polymers^42,43^.

## Conclusions

The focus of this work was the study of two reversing sequence amphiphilic synthetic peptides with a lipid like behavior at air/water interface. In the hydrophobic region, both peptides have an enriched tyrosine domain able to form di-tyrosine upon surface illumination. Both peptides are able to laterally conduct electrons almost 8 times higher than phospholipid. The conductivity of the confined peptide depends on the peptide bond backbone orientation being the **C_t_→N_t_** flow more efficient than the **N_t_→C_t_** ones. The di-Tyr formation increases the conductivity properties of both peptides, suggesting that the laterally formation of cross-linking tyrosines enhance lateral conductivity. When we measure the current-voltage dependence of each peptide under appropriate condition, we evaluate that both have semiconductor diode-like properties. We propose that this conductivity asymmetry may be of biological importance in transmembrane proteins given a novel conceptual framework at molecular level to explain the modulation of in-out information flow in the asymmetric biomembranes.

## Acknowledgements

Authors would like to thank to the support received from CONICET, ABA and GDF are fellowship and member of research career from this National Institution. We also thank grants and support from FONCYT (PICT 2016-1010), SECyT-UNC (Argentina) and from Aalborg University Hospital (Denmark).

## Materials and Methods

### Reagents and Chemicals

Water Milli Q Type II was used routinely in all measurements (Langmuir and conductivity). Peptides were purchased from Biomatik (www.biomatik.com, USA); peptides are higher purity better than 95 % by HPLC and checked by MS. DMSO and chloroform were from Merck (Darmstadt, Germany). DMPC lipids was from Avanti Polar lipids, Inc. (Alabaster, AL, USA).

### Monolayer formation and compression π-Area isotherms

Langmuir monolayers experiments were performed at a room constant temperature (25 ^o^C), essentially as indicated before^44^. Langmuir monolayers were formed on pure water using a 180 mL volume trough. We prepared stock solutions of about 10 mM of the peptides and DMPC lipid which were dissolved in pure DMSO and chloroform, respectively. Individual diluted working solutions were freshly prepared to ~ 1 μM. Around of 15-30 μl of individual working solution was directly spread on the surface of water. The isotherms were obtained by continuous compression of the automatic and simultaneous movement of both barriers that enclose the formed film. The compression rate used was setting at 10 mm^2^.min^−1^. Was used the Wilhelmy method for determining the surface pressure, putting a Pt plate on the surface of monolayer. The film area and surface pressure were continuously recorded using a software provided by KSV Instruments^®^ Ltd.

### Brewster Angle Microscopy

Peptide monolayers were spread on the surface area closed by the electrodes fixed on the interface and inside the illuminating area (see below). The films were directly observed by using a Brewster Angle Microscope (BAM) coupled to an EP3 Imaging Ellipsometer equipment (Accurion, Göttingen, Germany) with a 20x objective (Nikon, Japan, NA 0.35). The adequate amount of peptide was spread on the surface to achieve a high packing around 55 mN.m^−1^.

The imaging acquisition were sequentially taken each 10 min after UVB illumination, in order to detect change induced by UVB light on the peptide monolayer. Results obtained with BAM were compared with the spectrums obtained by using RAMAN Microscopy and Fluorescence Microscopy (FM).

### Illumination device coupled to BAM

The UV 265 nm lamp was fixed on the top of the trough of BAM equipment, thus the UV light hits on the monolayer. The laser BAM hits inside the centre of the circular illuminated area. The images from the monolayer were take each 10 min for the rest one hour.

### Heterogeneity measurement

Was take several (> 10) images from BAM equipment coupled to UVB illumination device after each illumination round. The illumination rounds were each 10 minutes until 1 hour for two peptide monolayer separately. The images were processed with FIJI-Image J software. We determine the heterogeneity from the brightness intensity of all the images obtained, comparing the standard deviation of grey values ratio to grey media values.

### Setting of electronic measurements

In both measurements (conductivity and diode behaviour) a more description of the in-house settings is given in Supplemental Material.

The monolayer was Illuminated using a UVB 265 nm lamp (Thorn Labs catalogue model M265L3). The surface electrodes were fixed on the interface and inside the illuminating area. Appropriate amount of peptide (dissolved in pure high quality DMSO) or lipid (dissolved in high quality chloroform) was spread on the surface area closed by the electrodes for each experimental condition in order to achieve a high packing condition (around 55 mN.m^−1^). The film was left by 3-5 min before illumination and/or conductivity measurement.

### Conductivity measurements

Conductivity measurement was performed in three different experimental conditions: *i)* without monolayer illumination and without polarization; *ii)* without monolayer illumination under polarization and *iii)* with monolayer illumination and under polarization (see Fig. 3b). Measurements were recorded during one hour using SerialPlot software simultaneously keeping constant the UV illumination in the third experimental condition described above. The circuit measures resistance then the values were transformed to conductivity values taking into account the electrode diameter and the distance of separation between them at the surface. See Supplemental material for further details.

### Diode behaviour measurements

The measurement was performed using two experimental conditions: *i)* without monolayer illumination and *ii)* after five-hour of monolayer illumination. Measurements were recorded during one hour using the PuTTY software. In this case our in-house circuit measures the current with the voltage variation from gate to drain and scan in the range from −8 V to 8 V. The current from gate to drain was predominant to the current from source to drain. See Supplemental material for further details.

## SUPPLEMENTARY MATERIAL

**Fig. S1.**
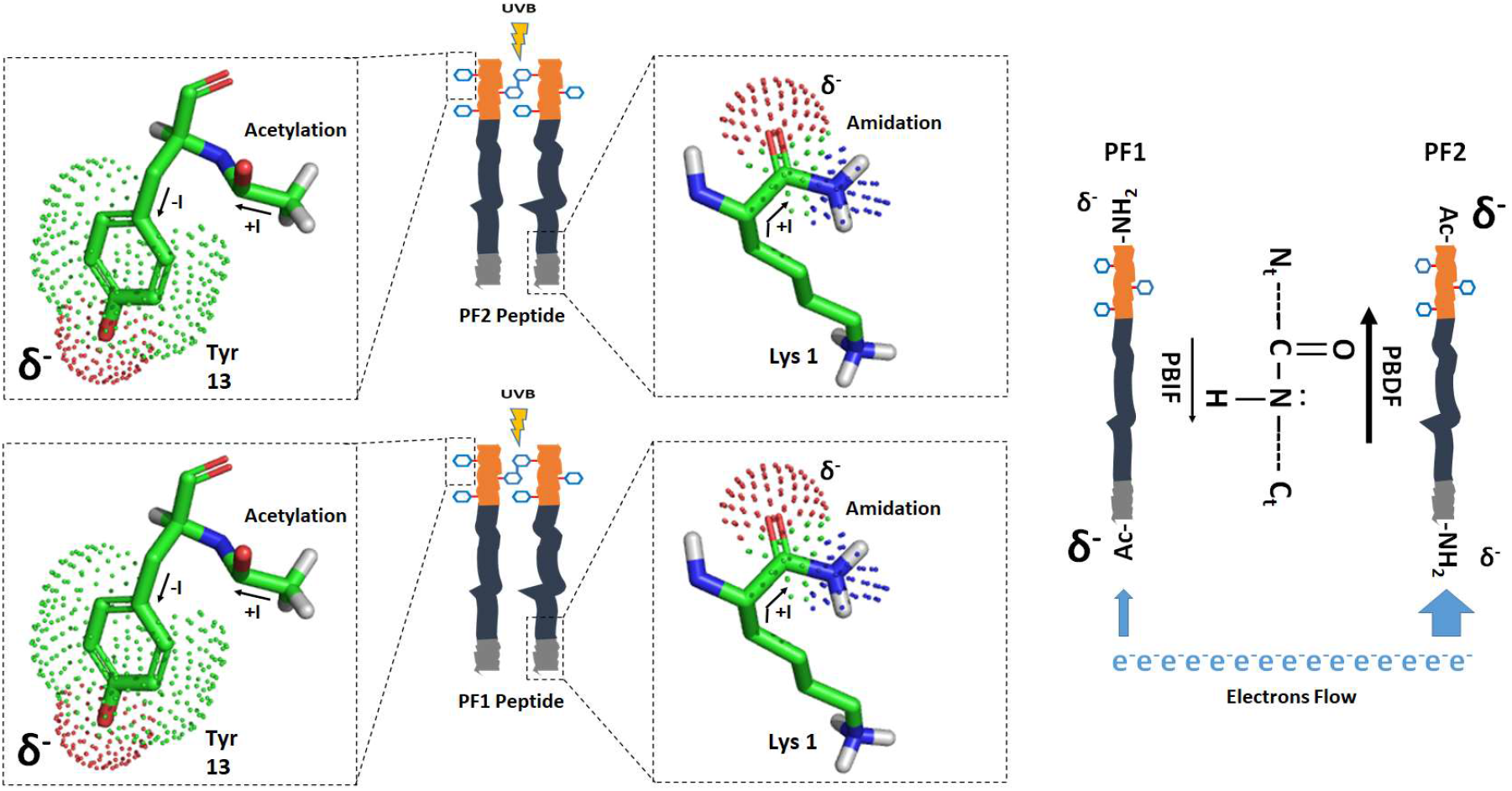
Influence of peptide acetylation and amidation on electron density. Schematic representation of the ending tyrosine (towards the air-end) and lysine residues (towards the water face) for both peptides. The bigger arrow in the right panel represents the higher conductivity throughout the peptide backbone of PF2 compared with PF1.

### Additional text to explain Fig S1

In a close packing array the more positive C-terminus region of PF2 is oriented towards to water end. This molecular configuration would offer less resistance to the electrons flow throughout the peptide backbone (see below). The fact that PF1 is originally acetylated but PF2 is amidated in the lysine which is in contact with the water phase would induce an overall less electropositive effect for PF2 compared to PF1 (i.e. more positive charge distribution oriented to the water subphase). This is because the PF1 C-N bound acetylation undergoes two inductive effect given by the side chain of lysine and, the other one from the −CH3 of the acetyl group. Instead, PF2 has only one inductive effect coming from the side chain of lysine. This description is a well-known effect in organic chemistry.^45^ This chemical description also explains an easier electron flow through the backbone for PF2 than PF1 (Fig. S1).

**Fig. S2.**
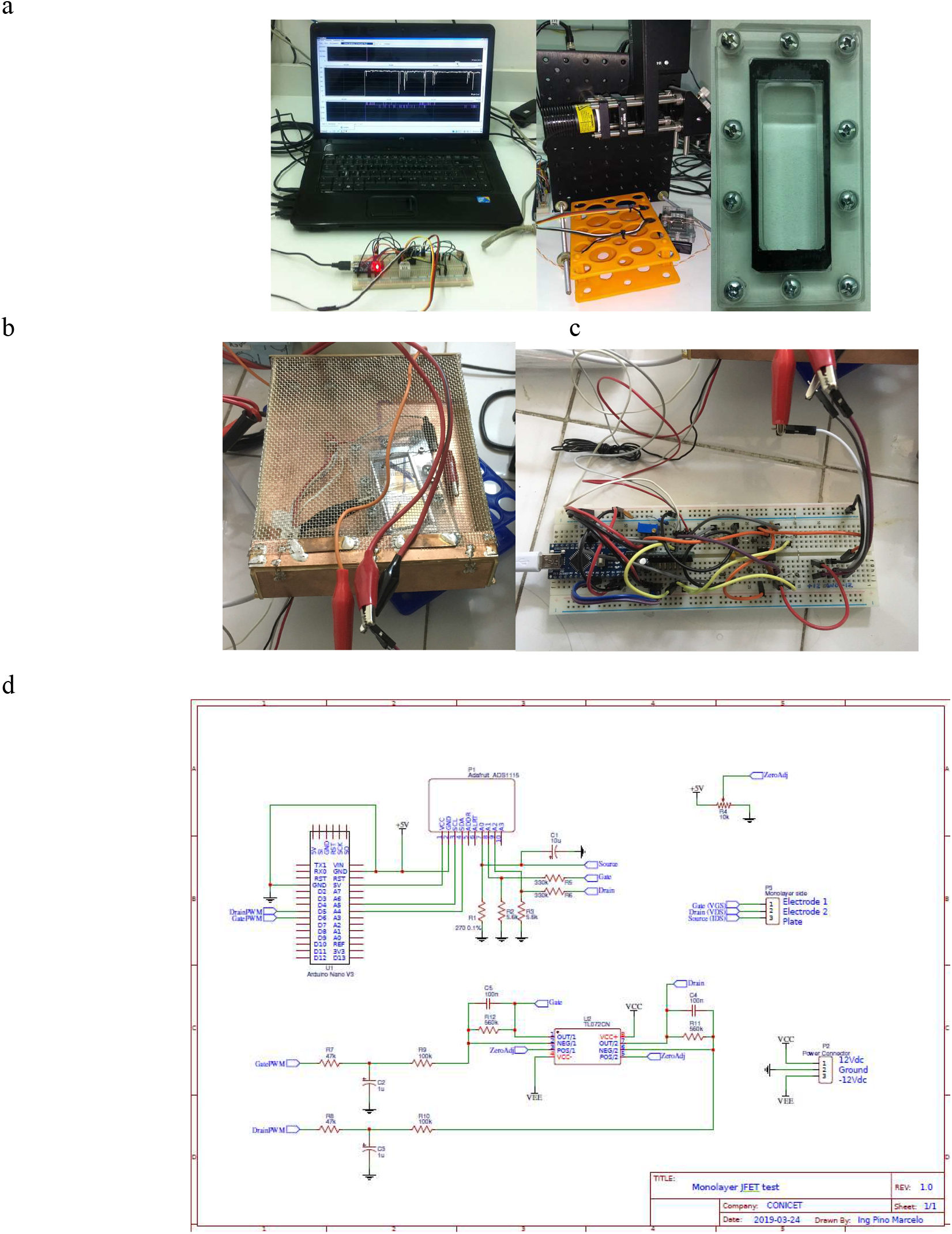
Setting devices for conductivity and voltage-current measurement and UVB illuminating source of peptide monolayers. **a**, Picture of electronic circuit connected to the PC, illuminating device with a 265 nm laser UVB source from Thorn Labs catalog model M265L3 (central panel) and the home-made Teflon mini trough for peptide spreading (right panel). The spreading surface area dimension are 75 mm wide x 300 mm length and 2 mm deep framed with an acrylic support. In the central part of the picture can be seen the Teflon trough with both surface electrodes for conductivity measurement. Each electrode was made of stainless-steel wire of 0.13 mm of diameter. **b**, Setting device for diode measurement. The trough is collocated inside of a metal Faraday cage (left panel); **c**, in-house electronic circuit couplet to PC computer. The polarization base of trough is a stainless-steel plate. **d**, electronic circuit diagram.

**Fig S3:**
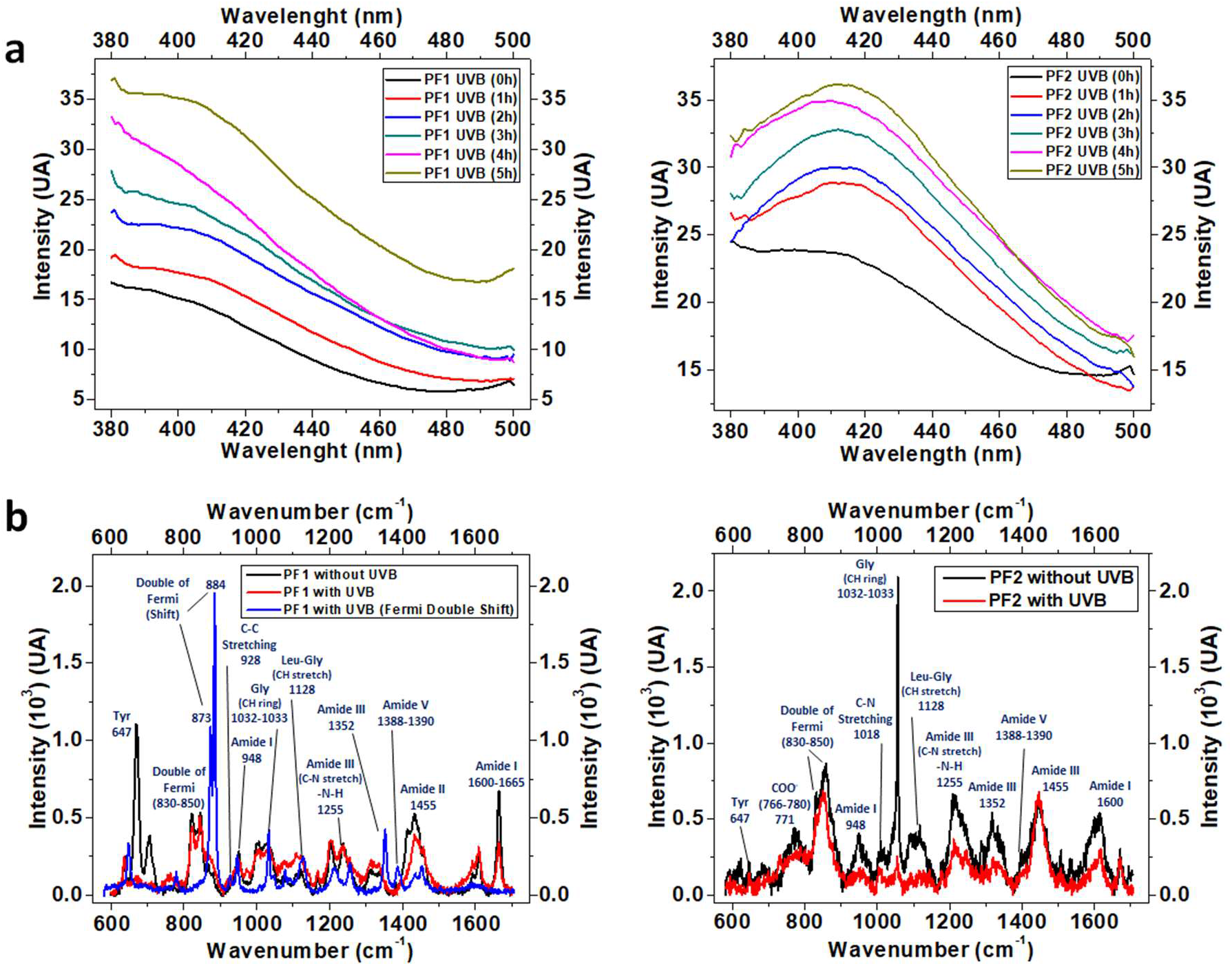
Di-Tyr identification of tyrosinated peptides. **a,** Fluorescence spectra of PF1 and PF2 peptides, due the di-tyr formation detected by the typical emission between 410 nm-420 nm. **b**, RAMAN spectra of PF1 and PF2 peptides. For PF2, after 10 min of UVB illumination at 265 nm, we detected double Fermi pick corresponding to tyrosine frequency vibration at the typical wavenumber values of 830 cm^−1^ and 850 cm^−1^. However, PF1 shows the typical shift of double Fermi pick to 873 cm^−1^ and 884 cm^−1^. The higher scattering observed for PF1 fluorescence (Fig S3a, left) is due to its greater heterogeneity observed for this peptide (see Fig. 2 of the paper) and it is keeping with changes in the double Fermi pick observed in RAMAN for this peptide (Fig. S3b, left).

### Additional text to Fig. S3

#### Surface formation of di-tyr of tyrosinated peptides upon UVB illumination

We have used three different laser lamps to illuminate the peptide monolayer either at 265, 280 or 300 nm as UVB source. We have observed the formation of di-tyr domains regarding of UVB source employed. The time detections were each 10 min for the first hour and each hour for the rest 5 hours (Fig. S3).

We have also used RAMAN microscopy to follow di-tyr formation. After 10 min of UVB illumination at 265nm we detected the Fermi double corresponding tyrosine frequency vibration at typical wavenumber values 830 cm^−1^ and 850 cm^−1^ for PF2^46^. However, PF1 shows a shift of typical Fermi double to 873 cm^−1^ and 884 cm^−1^, respectively, probably due to the formation of iso di-dyr similarly to the data reported previously^7,47^.

In addition, we detected an increase in the wavenumber range 1032 cm^−1^ to 1033 cm^−1^ for −CH ring of tyrosine in PF2 compared with PF1 at t_o_ (no UVB illumination). However, this wavenumber range is higher for PF1 than PF2 at t = 10 min. On other hand, we observed less changes in wavenumber frequencies in the range corresponding to Amide I, Amide II and Amide III, however, there are some changes in the relative intensity for post-illuminated films when compared with t_o_. This could be related to the differences observed in electron flow and conductivity through the peptide backbone.

#### Conductivity properties of tyrosinated peptides

**Fig. S4.**
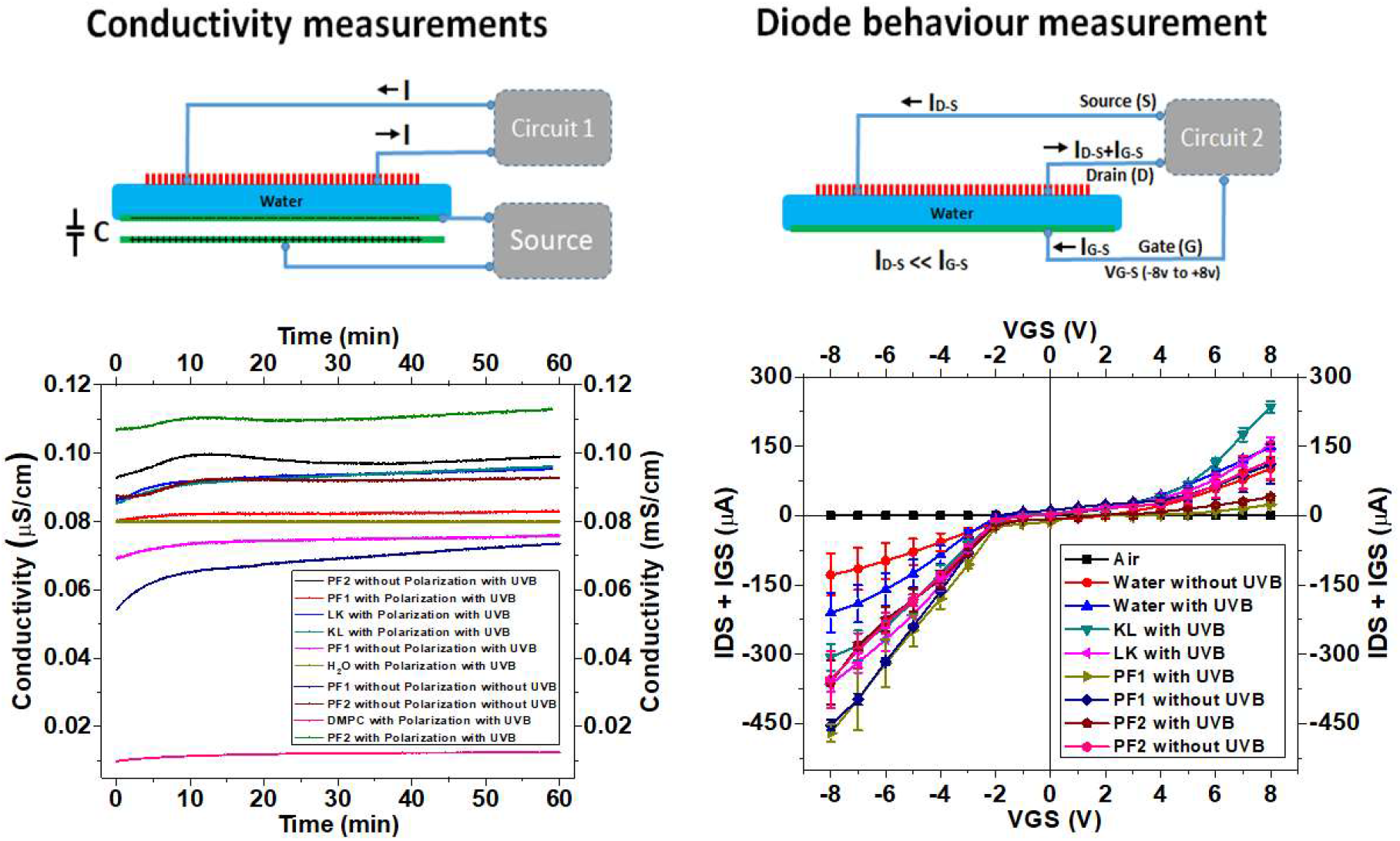
Electronic setting to conductivity and diode-like measurements in oriented tyrosine rich peptide monolayers. **Left panel:** Setting of conductivity measurements. Upper part: cartoon representation of the conductivity measurement setting. Lower part: graphic analysis of conductivity measurements. **Right panel:** Current-voltage dependence setting. Upper part: cartoon representation of the current-voltage dependence setting. Lower part: graphic analysis of current-voltage dependence. UVB: illumination at 265 nm when indicated for 1 h. Polarization with the metal plate source **on**. Sequence of non-tyrosinated peptides, KL: Ac-KKGWLLLLLL-NH_2_; LK: Ac-LLLLLLWGKK-NH_2_ (see reference^6^ for details about the surface properties of these peptides)

### Additional Text

#### Conceptual assimilation of peptide bond backbone to an inorganic semiconductor with diode behavior

Analogous to the general theory, peptides are based on carbon atom, located in the 4^th^ group of the periodic table similar to silicium and germanium atoms. Additionally, the peptide bond has nitrogen, located in the 5^th^ group of the periodic table (similar to the donor doping agent as arsenic or phosphorus atoms) and the oxygen, highly electronegativity, that can act as acceptor.^48^ However, oxygen is located in the 6^th^ group of the periodic table different to the acceptor doping agent such as barium or aluminium which are in the 3^ed^ group of the periodic table. It is well known that the peptide bond is a amide type covalent chemical bond stabilized by resonance^49^. Thus, oxygen pull electrons density due to its higher electronegativity acting as an electron acceptor dopant releasing electrons for conduction when an external electrical potential is applied. Hence, oxygen in the peptide bond network works as a temporally storage electrons. Contrary, nitrogen is an electron donor^28^ dopant that provides charge carriers which transfers electric current though the backbone help by the electronegativity of oxygen atom. Accordingly, nitrogen works as a temporally forming gaps and temporally storages electrons (Fig. 3d). With this vision in mine, we can conceptualize that if a single peptide bond acts as a diode, where nitrogen is the ***N*** material and oxygen is the ***P*** material, thus one polypeptide is a diode system connected in serial form (Fig. 3d). Taking into account what other authors have been suggested, regarding to the theory of electron tunnels for peptides,^50–52^ each peptide bond has a potential barrier to overcome the electrons flow from oxygen acceptor atom to the nitrogen atom as donor.

**Figure S5.**
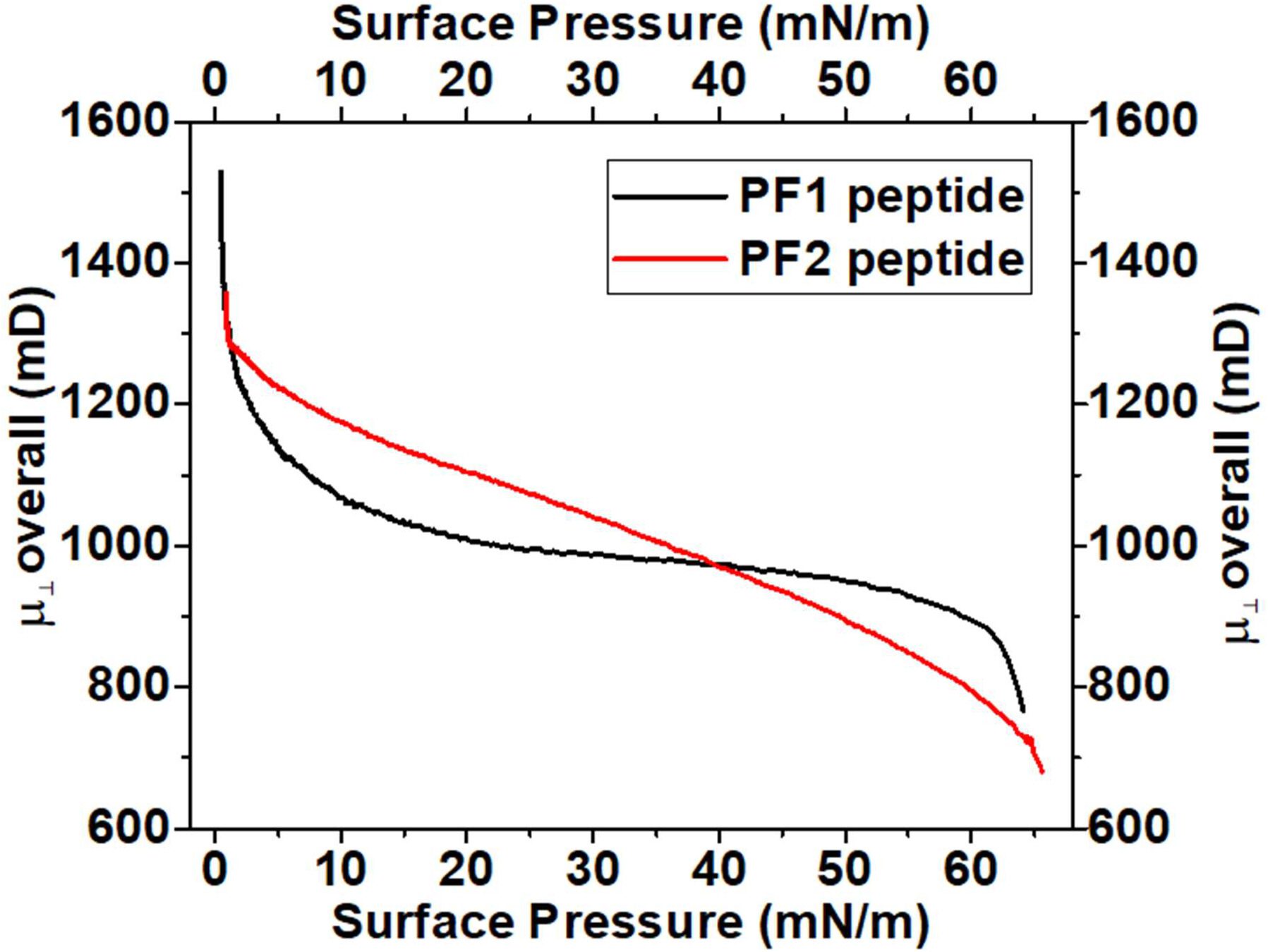
Individual Overall Surface Dipole Moment (μ_⊥_) for both peptides. The overall dipolar moment (μ_⊥_) towards the air end for an ionized monolayer (according to^20,21^) is given by: 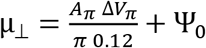 where *A_π_* and ΔV_*π*_ are the molecular area and the surface potential at a defined surface pressure *π*, in nm^2^ and millivolt units, respectively; Ψ_0_ is the potential difference of the ionic double layer of the ionized peptide film (Ψ_0_ values are practically the same because of the surface charge density is identical for both peptides). μ_⊥_ is the overall dipole moment perpendicular to the air/water interface resultant of three contribution according to μ_⊥_= μ_1_+ μ_2_ + μ_3_ (see^21,22^). μ_1_ comes from the oriented water molecules around the hydrophilic portion of the amphiphilic peptide; μ_2_ is the contribution of hydrophilic end corresponding to the lysine residues (equivalent to the polar head group in the lipids) and μ_3_ comes from the hydrophobic portion of the peptide corresponding to the leucine and tyrosine domains (equivalent to hydrocarbon chains in lipids), see Fig. 1 in the paper.

